# *MCOLN1* gene-replacement therapy corrects neurologic dysfunction in the mouse model of mucolipidosis IV

**DOI:** 10.1101/2020.12.06.413740

**Authors:** Samantha DeRosa, Monica Salani, Sierra Smith, Madison Sangster, Victoria Miller-Browne, Sarah Wassmer, Ru Xiao, Luk Vandenberghe, Susan Slaugenhaupt, Albert Misko, Yulia Grishchuk

## Abstract

Mucolipidosis IV (MLIV, OMIM 252650) is an orphan disease leading to debilitating psychomotor deficits and vision loss. It is caused by loss-of-function mutations in the *MCOLN1* gene that encodes thethe lysosomal transient receptor potential channel mucolipin 1 (TRPML1). With no existing therapy, the unmet need in this disease is very high. Here we show that AAV-mediated gene transfer of the human *MCOLN1* gene rescues motor function and alleviates brain pathology in the *Mcoln1^−/−^* MLIV mouse model. Using the AAV-PHP.b vector for initial proof-of-principle experiments in symptomatic mice, we showed long-term reversal of declined motor function and significant delay of paralysis. Next, we designed self-complimentary AAV9 vector for clinical use and showed that its intracerebroventricular administration in post-natal day 1 mice significantly improved motor function and myelination and reduced lysosomal storage load in the MLIV mouse brain. We also showed that CNS targeted gene transfer is necessary to achieve therapeutic efficacy in this disease. Based on our data and general advancements in the gene therapy field, we propose scAAV9-mediated CSF-targeted *MCOLN1* gene transfer as a therapeutic strategy in MLIV.

## Introduction

Mucolipidosis type IV (MLIV) is a lysosomal disorder caused by loss-of-function mutations in the *MCOLN1* gene (1). The clinical syndrome of MLIV was first described in 1974 (2) and the corresponding gene was reported in 1999 (3–5). Patients typically present in the first year of life with delayed developmental milestones and reach a plateau in psychomotor development by two years of age. Axial hypotonia and signs of pyramidal and extrapyramidal motor dysfunction manifest early in life, preventing independent ambulation and severely limiting fine motor function. Though MLIV was originally described as a static neurodevelopmental disorder, progressive neurological deterioration has been recently been documented. Patients frequently exhibit progressive spastic quadriplegia across the first decade of life with clear loss of gross and fine motor skills during the second decade (Dr. Misko, personal observations). In congruence with the clinical course, ancillary brain imaging has demonstrated stable white matter abnormalities (corpus callosum hypoplasia/dysgenesis and white matter lesions) with the emergence of subcortical volume loss and cerebellar atrophy in older patients. Visual impairment is also a prominent feature of MLIV with progressive retinal dystrophy and optic nerve atrophy leading to blindness by the second decade of life (6–9), further impeding function and negatively impacting quality of life. Currently, the standard of care for MLIV centers on symptomatic management and no disease modifying treatments are available.

*MCOLN1* encodes the late endosomal/lysosomal non-selective cationic ion channel TRPML1, that regulates lysosomal ion balance and is directly involved in multiple lysosome-related pathways, including Ca^2+^-mediated fusion/fission with lysosomal membrane, mTOR-signaling, TFEB activation, lysosomal biogenesis(10–13), and autophagosome formation (14). Additionally, its role in Fe^2+^-transport and regulation brain iron homeostasis has also been demonstrated (15, 16).

Important insights into the pathophysiology of MLIV have been obtained from our genetic mouse model, *Mcoln1* knock-out mouse (17–20). *Mcoln1^−/−^* mice recapitulate the clinical and pathological, phenotype of MLIV patients including motor deficits, retinal degeneration, and the primary pathological hallmarks of corpus callosum hypoplasia, microgliosis, astrocytosis and, late, partial loss of Purkinje cells.

In this study, we demonstrated that *MCOLN1* gene transfer using AAV-PHP.b can either prevent or fully reverse neurological dysfunction in the MLIV mouse model, depending on the timing of treatment. To strengthen the translational potential of this study we next designed and tested a self-complimentary AAV9-based approach which showed a similar therapeutic efficacy that was dependent on adequate brain transduction. Altogether, this pre-clinical study sets a stage for developing AAV-mediated CNS-targeted gene transfer as a therapeutic modality for this devastating and currently untreatable disease.

## Results

### *MCOLN1* gene transfer in juvenile pre-symptomatic *Mcoln1^−/−^* mice prevents onset of motor deficits

*Mcoln1^−/−^* mice mimic all the primary manifestations of the human mucolipidosis IV disease, including neurologic deficits, brain pathology, gastric parietal cell dysfunction with high systemic gastrin, and retinal degeneration (16–22). The earliest motor deficits appear in the form of reduced vertical activity in the open field test at the age of two months **(Suppl. Fig 1)**. Motor dysfunction in *Mcoln1^−/−^* mice is progressive and results in decreased performance in the rotarod test at four months of age **(Figures 2 C,D and 6 D, E)**, significant gait deficits (18) and, eventually, hind limb paralysis and pre-mature death at 7-8 months of age (18). Here we tested whether CNS-targeted transfer of the human *MCOLN1* gene in juvenile pre-symptomatic KO mice (5-6 weeks of age) prevents motor deficits usually occurring at 2 months of age. We intravenously administered either saline or AAV-PHP.b-CMV-*MCOLN1*-F[urin]F2a-eGFP (later in text referred to as PHP.b-*MCOLN1*) intravenously to *Mcoln1^−/−^* and control *Mcoln1^−/+^* male mice at the age of 5-6 weeks after they reached a body mass of 19 g. Four weeks after injections, at 2 months of age, mice were tested in the open field arena and euthanized for tissue collection **(Table 1)**. *Mcoln1^−/−^* mice treated with PHP.b-*MCOLN1* showed normal vertical activity unlike saline-treated *Mcoln1^−/−^* mice which demonstrated the expected decline **(Figure 1A, B)**. Post-mortem tissue analysis showed high expression of the human *MCOLN1* transgene in the brain parenchyma, particularly, in the cerebral cortex and cerebellum, and detectable expression in peripheral tissues such as liver and muscle **(Figure 1C)**. Of note, the human-specific *MCOLN1* Taq-man assay that we used for transcriptional analysis (Hs01100653_m1, Applied Biosystems) has 87-94% homology with the mouse *Mcoln1* sequence and detects endogenous murine *Mcoln1* transcripts in a dose-specific manner as demonstrated on the graphs in **Figure 1C** for *Mcoln1^+/+^*, *Mcoln1^−/+^* and *Mcoln1^−/−^* saline-treated samples.

**Table 1.** All vector and trial summary.

| **Cohort** | **Treatment group** | **N** | **RO A** | **Dose** | **Age at treatment** | **Age at takedown** |
| --- | --- | --- | --- | --- | --- | --- |
| **1** | **WT-SALINE** | 9 | IV | 1e12 vg/mouse | 5-6 weeks (pre-symptomatic) | 9-10 weeks |
|  | **HET-SALINE** | 18 |  |  |  |  |
|  | **KO-SALINE** | 9 |  |  |  |  |
|  | **KO-AAV-PHP.b-MCOLN1** | 14 |  |  |  |  |
| **2** | **HET- SALINE** | 15 | IV | 1e12 vg/mouse | 2 months (symptomatic) | 11 months (around 7 months for untreated KO) |
|  | **KO-SALINE** | 12 |  |  |  |  |
|  | **KO-AAV-PHP.b-MCOLN1** | 9 |  |  |  |  |
| **3** | **HET-SALINE** | 20 | ICV |  | P1 (neonatal) | 9-10 weeks |
|  | **KO-SALINE** | 10 |  |  |  |  |
|  | **KO-scAAV9-JeT-MCOLN1** | 7 |  | 2e10 vg/mouse |  |  |
|  | **KO-scAAV9-JeT-MCOLN1** | 16 |  | 1e10 vg/mouse |  |  |
|  | **KO-scAAV9-JeT-MCOLN1** | 10 |  | 4e9 vg/mouse |  |  |
|  | **KO-scAAV9-SYN1-MCOLN1** | 10 |  | 2e10 vg/mouse |  |  |
| **4** | **HET- SALINE** | 16 | IV | 5e11 vg/mouse | 2 months (symptomatic) | 7 months |
|  | **KO-SALINE** | 10 |  |  |  |  |
|  | **KO-scAAV9-JeT-MCOLN1** | 9 |  |  |  |  |

**Figure 1.**
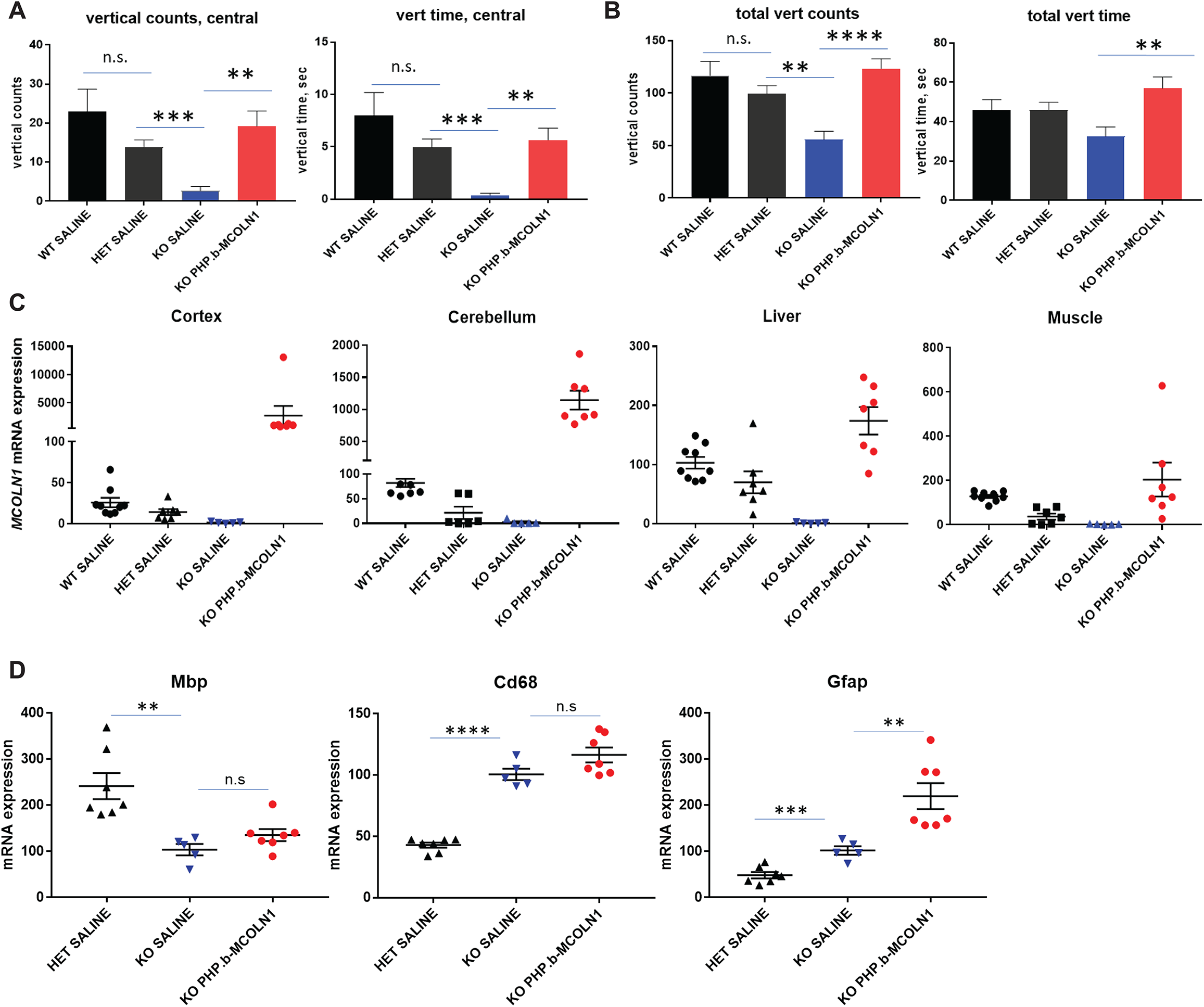
**Pre-symptomatic administration of AAV-PHP.b-*MCOLN1* prevents onset of early motor deficits in *Mcoln1^−/−^*mice. A, B.** Vertical activity measurements in the open field test, represented as vertical counts and vertical time in either central (A) or total area (B) of the open field arena, show significant impairment in the saline-treated 2 months-old *Mcoln1^−/−^* mice that was fully prevented in the *Mcoln1^−/−^* mice treated with AAV-PHP.b-*MCOLN1* intravenously at the age of 5-6 weeks; n (WT SALINE)=9; n (HET SALINE)=18; n (KO SALINE)=9; n (KO PHP.b-*MCOLN1*)=14. Data presented as mean values and SEM; group comparisons made using unpaired T-test. **C.** qRT-PCR analysis of the *MCOLN1* transcripts showing transgene overexpression in the cerebral cortex, cerebellum, liver and quad muscle. Data presented as mean values and SEM; n (WT SALINE)=9; n (HET SALINE)=6-7; n (KO SALINE)=5; n (KO PHP.b-*MCOLN1*)=7. **D.** qRT-PCR analysis of the myelination marker *Mbp*, microgliosis marker *Cd68* and astrocytosis marker *Gfap* in the cerebral cortex; n (HET SALINE)=7; n (KO SALINE)=5; n (KO PHP.b-*MCOLN1*)=7. Data presented as mean values and SEM, group comparisons were made using un-paired T-test.

The major histopathological hallmarks in the brain *Mcoln1^−/−^* mice at 2 months of age include reduced brain myelination, microgliosis and astrocytosis (16, 20, 21, 23). Despite the strong effect of the *MCOLN1* gene transfer on motor function in this cohort **(Figure 1A)**, qRT-PCR analysis of the post-mortem brain tissue showed no changes in expression of the myelination marker *Mbp* and microglial activation marker *Cd68* **(Figure 1D)**. Unexpectedly, we observed elevated expression of the astrocyte activation marker *Gfap* in PHP.b-*MCOLN1*-treated *Mcoln1^−/−^* mice compared to saline-treated *Mcoln1^−/−^* and *Mcoln1^+/−^* groups. Increased astrocyte activation in treated mice may reflect a response to the viral vector or expression of a non-self protein. Importantly, we observed no overt health concerns in treated mice during 4 weeks of post-treatment observation.

### *MCOLN1* gene transfer in symptomatic *Mcoln1^−/−^* mice restores motor function and delays onset of paralysis

In the first years of life, MLIV patients exhibit severe impairment of gross and fine motor development. Patient care givers identify developmental motor dysfunction as a major contributor to neurological disability and limited quality of life. Taking this into account, a successful treatment for MLIV should aim to rescue developmental motor deficits rather than delay the later onset of motor deterioration alone. To test whether *MCOLN1* transgene expression *restores* motor function *after* symptom onset, *Mcoln1^−/−^* mice were intravenously administered either saline or PHP.b-*MCOLN1* at two months of age **(Table 1)**. Motor function was assessed in the open field test once at the age of 4 months and, subsequently, by monthly rotarod testing until the completion of the study. Open field revealed significant rescue of vertical activity in the PHP.b-*MCOLN1* – treated *Mcoln1^−/−^* mice when analyzed in either the central zone **(Figure 2A)** or in the entire area of the arena **(Figure 2B),** demonstrating for the first time that therapeutic delivery of the *MCOLN1* gene can restore neurologic function in MLIV, even after it has already been compromised. In rotarod testing, saline-treated *Mcoln1^−/−^* mice showed a higher probability to fall at the age of 4 and 5 months **(Figure 2C)** and lower average latency to fall **(Figure 2D).**

At the age of 6 months, saline-treated *Mcoln1^−/−^* mice developed profound hind limb weakness and were excluded from testing. By 7 months of age, they developed hind-limb paralysis and were euthanized in accordance with humane criteria **(Figure 2E).** Remarkably, PHP.b-*MCOLN1*-treated *Mcoln1^−/−^* mice were indistinguishable from the control healthy littermates in the rotarod test until the end of the trial at 11 months of age **(Figure 2 C, D).** Only two out of nine PHP.b-*MCOLN1*-treated *Mcoln1^−/−^* mice showed signs of paralysis after 8 months of age. All other *Mcoln1^−/−^*-treated mice survived over 4 months longer than untreated controls **(Figure 2E)** and showed no signs of functional decline at the time of euthanasia at 11 months of age.

**Figure 2.**
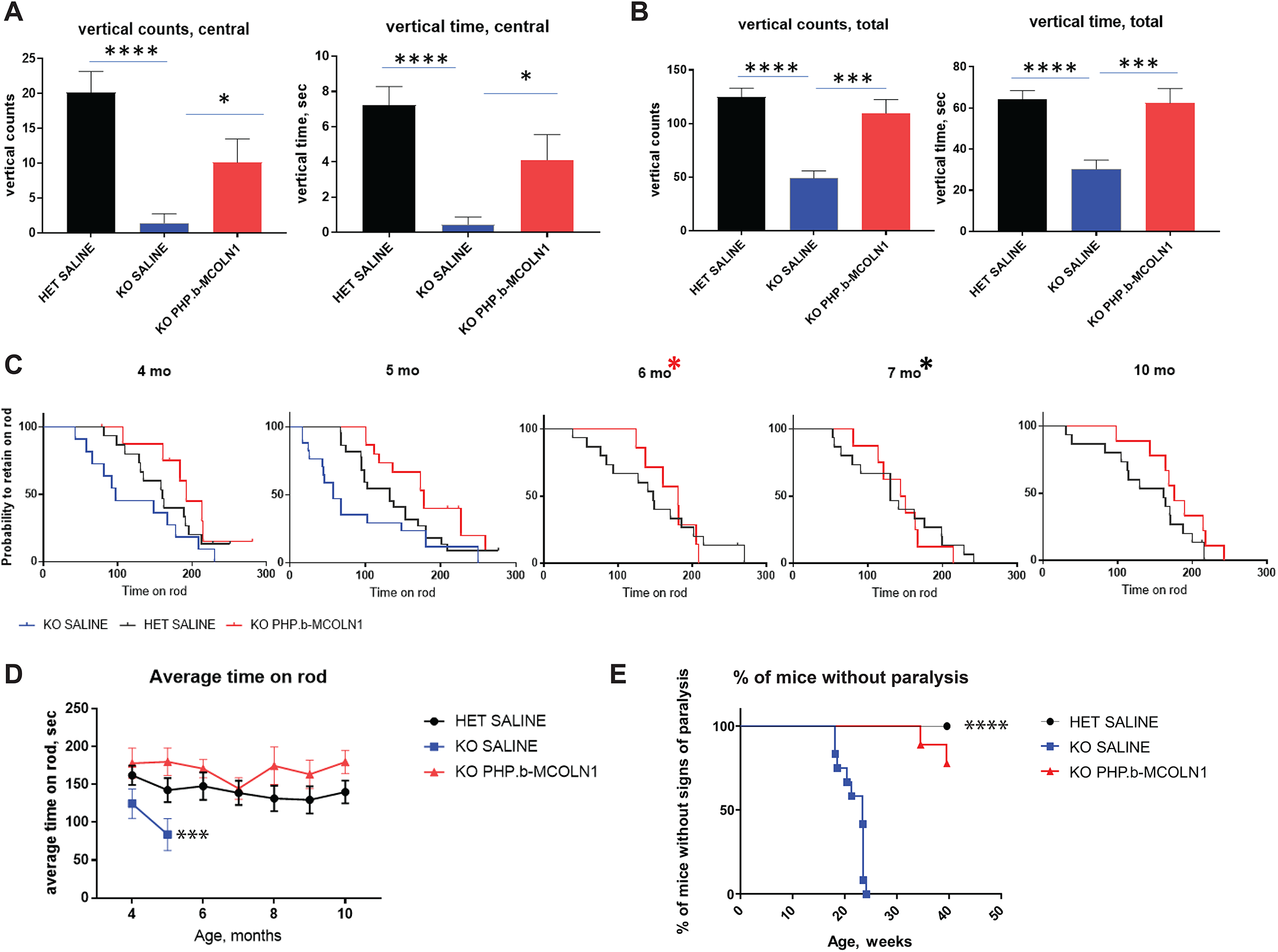
**Systemic administration of AAV-PHP.b-*MCOLN1* in symptomatic *Mcoln1^−/−^* mice restores motor function and significantly extends time to paralysis. A, B.** Measurements of the vertical activity in the open field test, represented as vertical counts and vertical time in either central (A) or total area (B) of the open field arena, show significantly impaired activity in the 4 months-old *Mcoln1^−/−^* mice treated with saline, and its significant recovery in the *Mcoln1^−/−^* mice that were treated with AAV-PHP.b-*MCOLN1* at the symptomatic stage of the disease at 2 months of age; n (HET SALINE)=15; n (KO SALINE)=12; n (KO PHP.b-*MCOLN1*)=9. Data presented as mean values and SEM; group comparisons made using un-paired T-test. **C, D.** Significant long-term improvement of the rotarod performance presented as a probability to retain on the rod **(C)** or average latency to fall **(D)** in the *Mcoln1^−/−^* mice treated with AAV-PHP.b-*MCOLN1.* **C.** Log-rank analysis of the probability-to-retain-on-the-rod curves revealed significant changes between curves, at the age of 4 (p=0.036) and 5 months (p=0.0052) due to worse rotarod performance of the KO SALINE, group, and no changes between HET SALINE and KO PHP.b-*MCOLN1* curves at 6, 7, 8, 9, 10 months (see also supplementary figure 2). Red asterisk at 6 mo marks the time point when KO SALINE mice developed hind-limbs weakness and had to be excluded from testing; black asterisk at 7 mo marks the time-point when all the KO SALINE mice were euthanized due to paralysis; n (HET SALINE)=14; n (KO SALINE)=12; n (KO PHP.b-*MCOLN1*)=9. **D.** Two-way ANOVA test (treatment group × age) of the average latency to fall for KO SALINE and KO PHP.b-*MCOLN1* groups at 4 and 5 mo time points shows significant effect of AAV treatment, p=0.0007; F=13.62. **E.** Systemic administration of AAV PHP.b-*MCOLN1* significantly delays time to paralysis in *Mcoln1^−/−^* mice. The criterion for paralysis was loss of righting reflex when mouse failed to rotate itself in upright position after placing on a side within 10 sec; n (HET SALINE)=14; n (KO SALINE)=12; n (KO PHP.b-*MCOLN1*)=9; log-rank test p-value is <0.0001.

qPCR analysis showed long-term (tissues were collected 9 months after vector administration) human *MCOLN1* transgene expression in the brain tissue, specifically in the cerebral cortex and cerebellum, and in peripheral organs, such as liver and muscle. Interestingly, we also measured high *MCOLN1* transgene expression in the sciatic nerves, indicating transduction of peripheral nerves **(Figure 3A)**.

**Figure 3.**
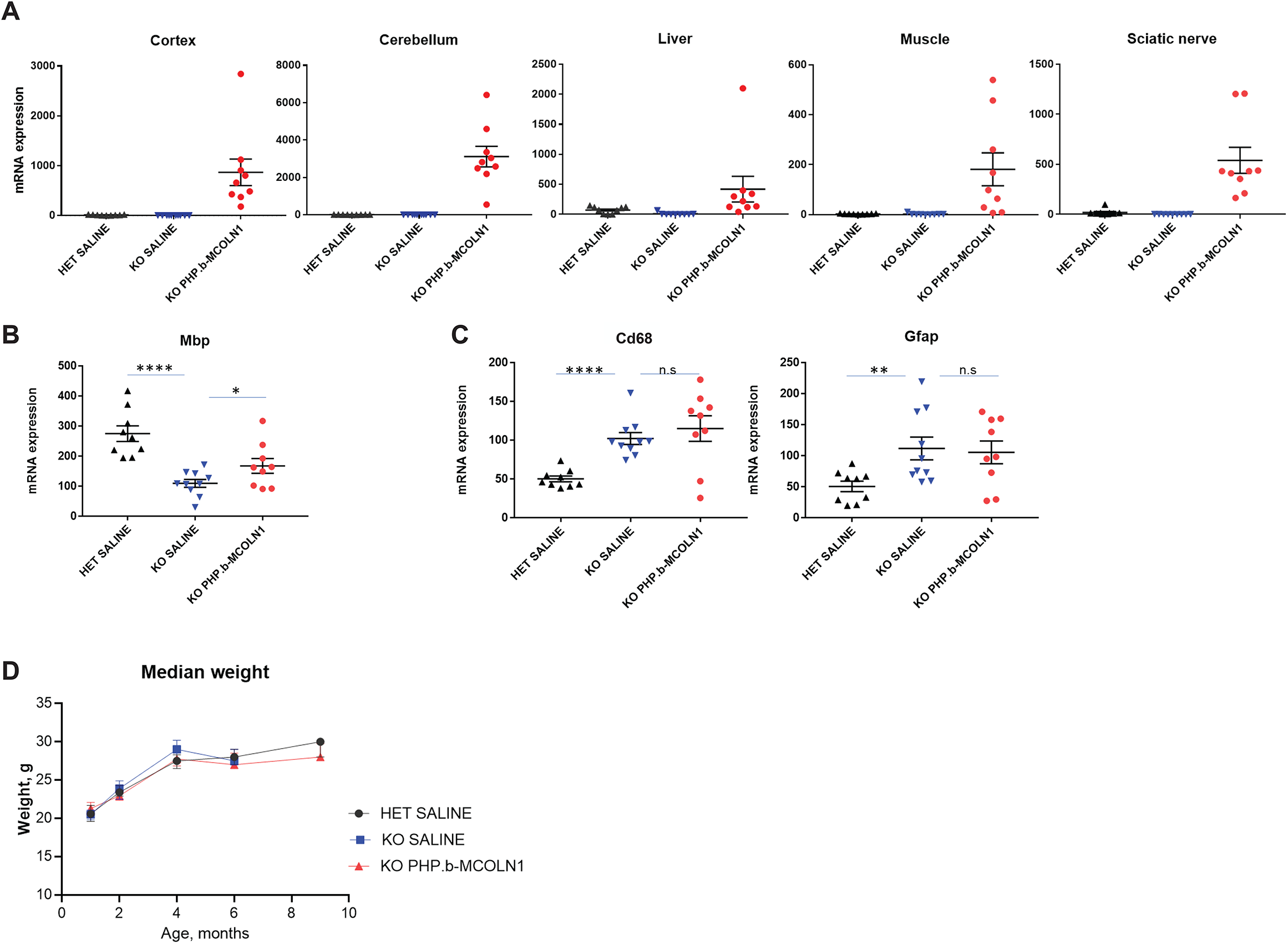
**Biodistribution and efficacy of systemic administration of AAV-PHP.b-*MCOLN1* in symptomatic *Mcoln1^−/−^* mice. A.** qRT-PCR analysis of the *MCOLN1* transcripts showing transgene overexpression in the cerebral cortex, cerebellum, liver, quad muscle and sciatic nerve. Data presented as mean values and SEM; n (HET SALINE)=14; n (KO SALINE)=12; n (KO PHP.b-*MCOLN1*)=9. **B.** qRT-PCR analysis of the myelination marker *Mbp* in cerebral cortex shows partial recovery in the PHP.b-*MCOLN1-* treated *Mcoln1^−/−^* mice. Data presented as mean values and SEM; n (HET SALINE) =14; n (KO SALINE) =12; n (KO PHP.b-*MCOLN1*) =9; group comparisons were made using unpaired T-test. **C.** qRT-PCR analysis of the microgliosis marker *Cd68* and astrocytosis marker *Gfap* in the cerebral cortex shows increased glial activation in the sham-treated *Mcoln1^−/−^* mice that was not changed in the PHP.b-*MCOLN1-* treated *Mcoln1^−/−^* mice; n (HET SALINE) =14; n (KO SALINE) =12; n (KO PHP.b-*MCOLN1*) =9. Data presented as mean values and SEM, group comparisons were made using unpaired T-test. **D.** No significant weight changes have been observed in either experimental group. Data presented as median values and interquartile range, n (HET SALINE) =14; n (KO SALINE) =12; n (KO PHP.b-*MCOLN1*) =9.

Administration of the PHP.b-MCOLN1 vector to *Mcoln1^−/−^* mice at two months of age resulted in partial but statistically significant rescue of brain myelination marker *Mbp* **(Figure 3B)** in the cerebral cortex demonstrating that myelination deficits in MLIV can be rescued even after the developmental course of brain myelination is complete. Notably, despite robust rescue of the motor function and high expression in the brain tissue, we observed no changes in the expression of the astrocytosis and microgliosis markers, *Gfap* and *Cd68*, in the cerebral cortex of PHP.b-*MCOLN1*-treated vs. saline-treated *Mcoln1^−/−^* mice **(Figure 3C)**. Administration of PHP.b-*MCOLN1* did not lead to any noticeable adverse effects, and no significant changes in weight have been observed between *Mcoln1^−/−^*-treated, *Mcoln1^−/−^* vehicle-treated and *Mcoln1^−/+^* vehicle-treated control groups **(Figure 3D)**.

### Intracerebroventricular administration of self-complementary AAV9-JeT-*MCOLN1* is efficacious in the *Mcoln1^−/−^* mouse model of MLIV

While AAV-PHP.b shows high brain transduction with systemic administration, its ability to penetrate blood-brain barrier is species-specific with the highest transduction rate in the C57Bl6 mice and a very low brain transduction in non-human primates (NHP) (24, 25). Therefore, we designed an *MCOLN1* expression vector suitable for gene transfer in human patients based on the clinically tested self-complimentary AAV9 vector (26–28). To drive expression of *MCOLN1,* we selected the synthetic promoter JeT, which is currently being used in a clinical trial of GAN gene transfer for giant axonal neuropathy (26, 29). For the most efficient and broad targeting of the CNS with scAAV9 we performed intracerebroventricular (ICV) injections in newborn mice at postnatal day 1. At birth, litters were randomly assigned to treatment with either saline or one of the three doses of the scAAV9-JeT-*MCOLN1* vector (2×10^10^ 1×10^10^ and 4×10^9^ vg/mouse). Injected mice were weaned and genotyped according to the standard procedure and subjected to open-field testing at the age of two months followed by euthanasia for tissue collection. qRT-PCR analysis revealed dose-dependent expression of the *MCOLN1* transgene in the brain and peripheral tissues, with the highest expression in the cerebral cortex **(Figure 4A)**. ICV administration of 2×10^10^) vg/mouse of scAAV9-JeT-*MCOLN1* (the highest dose used) prevented decline of motor deficits in the *Mcoln1^−/−^* mice at 2 months of age **(Figure 4 B, C)**. Post-mortem histological analysis of the brain tissue showed significant reduction in the density of lysosomal aggregates in the *Mcoln1^−/−^* cerebral cortex as demonstrated on representative LAMP1 immunostaining images **(Figure 4D)** and their quantification **(Figure 4E).** LAMP1-positive lysosomal aggregates are a prominent pathological feature of the *Mcoln1^−/−^* brain that can be quantified either by size of LAMP1-positive particles or as a percentage of the LAMP1-positive area. Both measures were reduced by *MCOLN1* gene transfer in the scAAV9-JeT-*MCOLN1-*treated *Mcoln1^−/−^* mice **(Figure 4E)**. Importantly, histological analysis of brain myelination using immunostaining for the myelination marker *Mbp* showed, an increased *Mbp-*positive corpus callosum area in the scAAV9-JeT-*MCOLN1-*treated *Mcoln1^−/−^* mice **(Figure 4F)**. Since developmental agenesis of the corpus callosum is a prominent feature of the human MLIV brain pathology, this data may have important implications for clinical trials measuring treatment efficacy in patients. Using qRT-PCR analysis in the cortical tissue we did not observe a significant increase in the expression of *Mbp* with any dose of scAAV9-JeT-*MCOLN1* **(Supplementary figure 3A)**, showing that recovery of myelination is primarily seen in the white matter. Additionally, while we noted a trend toward attenuation of microglial and astrocyte activation in the *Mcoln1^−/−^* cortical tissue in mice treated with scAAV9-JeT-*MCOLN1* using qRT-PCR analysis for a microglial and astrocyte activation markers *Cd68* and *Gfap*, the difference between *Mcoln1^−/−^*-saline- and scAAV9-JeT-*MCOLN1*-treated groups did not reach statistical significance with any of the vector doses used **(Supplementary figure 3A).**

**Figure 4.**
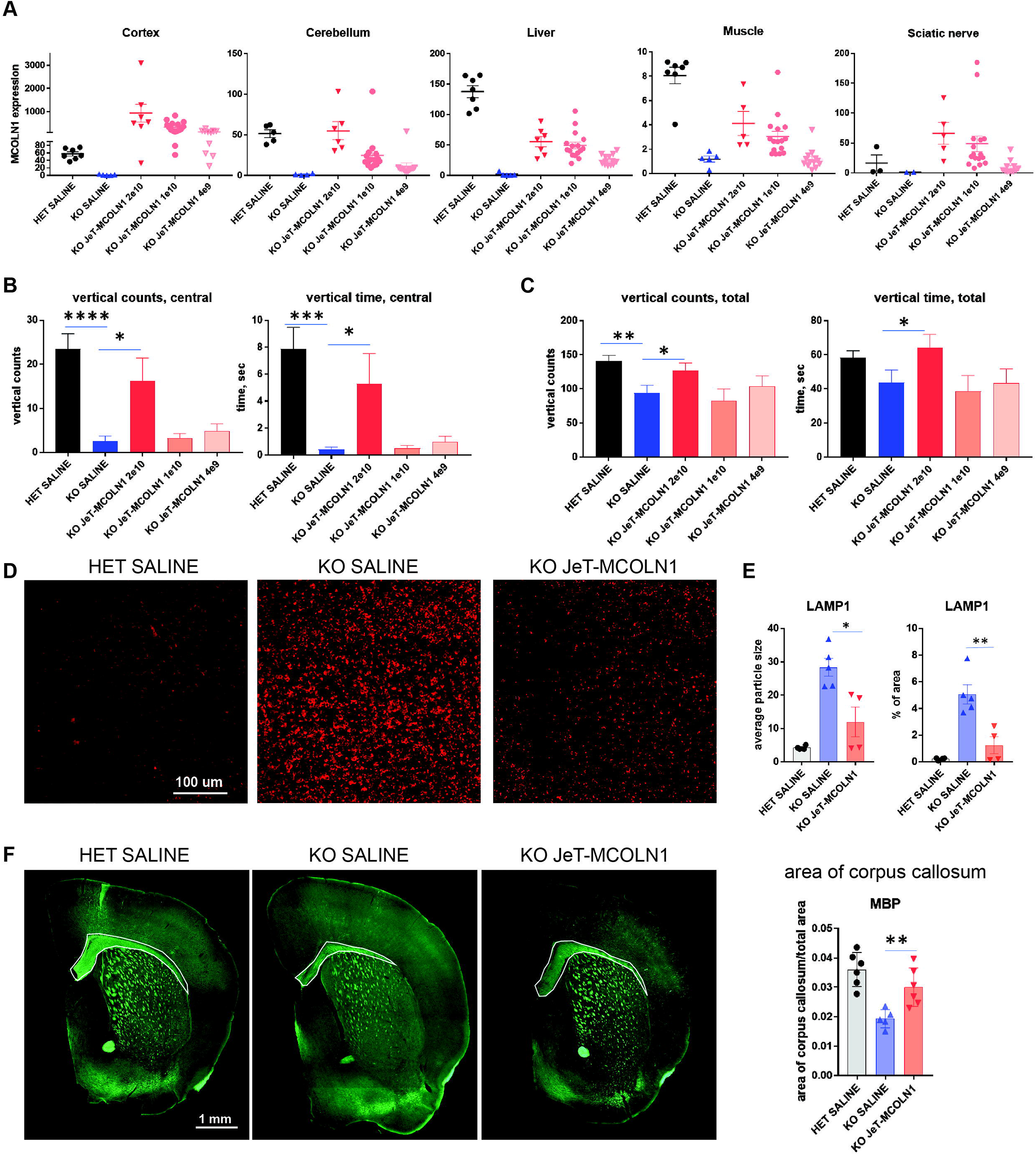
**The biodistribution and efficacy of intracerebroventricular delivery of scAAV9-JeT-*MCOLN1* in *Mcoln1^−/−^* neonatal mice. A.** qRT-PCR analysis of the *MCOLN1* transcripts demonstrates dose-dependent expression of *MCOLN1* in cerebral cortex, cerebellum, liver, quad muscle, and sciatic nerve. Data presented as mean values and SEM; n (HET SALINE)=3-7; n (KO SALINE)= 2-5; n (KO scAAV9-JeT-*MCOLN1;* 2e10 vg/mouse)=5-7; n (KO scAAV9-JeT-*MCOLN1;* 1e10 vg/mouse)= 16; n (KO scAAV9-JeT-*MCOLN1;* 4e9 vg/mouse)= 12. **B, C.** Vertical activity measured in the open field test as vertical counts and vertical time in either central **(B)** or total area **(C)** of the open field arena is significantly improved in *Mcoln1^−/−^* mice treated with the highest dose of scAAV9-JeT-*MCOLN1* (2e10 vg/mouse) injected ICV on post-natal day 1. Data presented as mean values and SEM; n (HET SALINE)=20; n (KO SALINE)= 10; n (KO scAAV9-JeT-*MCOLN1;* 2e10 vg/mouse)=7; n (KO scAAV9-JeT-*MCOLN1;* 1e10 vg/mouse)= 5; n (KO scAAV9-JeT-*MCOLN1;* 4e9 vg/mouse)= 10; group comparisons made using unpaired T-test. **D.** ICV delivery of scAAV9-JeT-*MCOLN1* reduces LAMP-positive lysosomal aggregates in the brain of *Mcoln1^−/−^* mice as demonstrated by LAMP immunohistochemistry representative images and blinded quantitative analysis on the right. LAMP staining presented as average particle size and percent of LAMP-positive area. Data presented as mean values and SEM, group comparisons made using unpaired T-test; n (HET SALINE)=4; n (KO SALINE)= 5; n (KO scAAV9-JeT-*MCOLN1;* 2e10 vg/mouse)=4. **E.** ICV delivery of scAAV9-JeT-*MCOLN1* improves corpus callosum thickness in the *Mcoln1^−/−^* mice. Representative images of MBP staining in the left panel show thicker corpus callosum (white selection). Blinded quantitative image analysis (right panel) shows enlarged area of the MBP-positive corpus callosum. Data presented as mean values and SEM, group comparisons made using unpaired T-test; n (HET SALINE)=6; n (KO SALINE)= 5; n (KO scAAV9-JeT-*MCOLN1;* 2e10 vg/mouse)=5.

### Neuron-specific expression of *MCOLN1* is sufficient to rescue early motor dysfunction in MLIV mice

Neonatal intracerebroventricular delivery of scAAV9 vectors results in the widespread transduction of neurons and a smaller fraction of glial cells in the mouse brain (30, 31), and leads to transduction of peripheral organs (31). To determine whether neuron-specific gene transfer of *MCOLN1* is responsible for the therapeutic recovery of neurological function in the *Mcoln1^−/−^* mouse model we next replaced the ubiquitously expressed JeT promoter with the neuron-specific human synapsin 1 (SYN1) promoter (31, 32). We administered the scAAV9-SYN1-*MCOLN1* vector to the postnatal day 1 mice via ICV route at the highest dose we used in the scAAV9-JeT-*MCOLN1* experiment, 2×10^10^ vg/mouse, for direct comparison of the two vectors **(Table 1)**. scAAV9-SYN1-*MCOLN1* – treated *Mcoln1^−/−^* mice demonstrated motor function recovery similar to the scAAV9-JeT-MCOLN1-treated *Mcoln1^−/−^* group **(Figure 5A, B)**.

**Figure 5.**
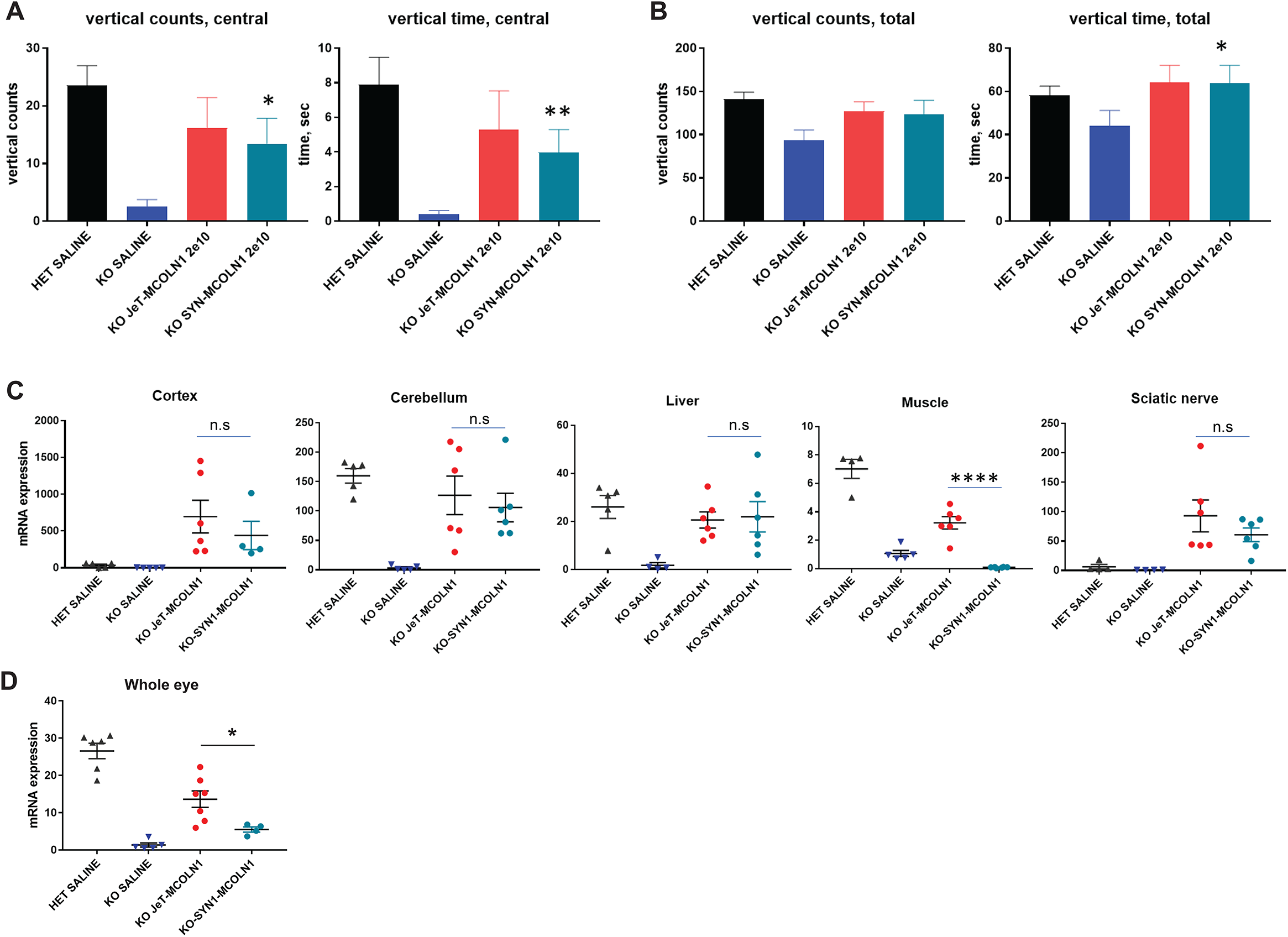
**Neuron-specific expression of *MCOLN1* is sufficient to rescue early motor dysfunction in *Mcoln1^−/−^* mice. A, B.** Vertical activity, measured in the open field test as vertical counts and vertical time in either central **(A)** or total area **(B)** of the open field arena, is significantly improved in *Mcoln1^−/−^* mice, treated with the scAAV9-SYN1-*MCOLN1* (neuron-specific expression) similarly to scAAV9-JeT-*MCOLN1* (ubiquitous expression). Both vectors were injected ICV at 2×10 vg/mouse on post-natal day 1. Data presented as mean values and SEM; n (HET SALINE)=20; n (KO SALINE)= 10; n (KO scAAV9-JeT-*MCOLN1;* 2×10 vg/mouse)=7; n (KO scAAV9-SYN1-*MCOLN1;* 2×10 vg/mouse)= 10; group comparisons made using unpaired T-test. **C.** qRT-PCR analysis of the *MCOLN1* transcripts shows expression of the transgene in cerebral cortex, cerebellum, liver, quad muscle, and sciatic nerve. Data presented as mean values and SEM; n (HET SALINE)=4-5; n (KO SALINE)= 4-5; n (KO scAAV9-JeT-*MCOLN1;* 2e10 vg/mouse)=6; n (KO scAAV9-SYN1-*MCOLN1;* 2e10 vg/mouse)= 4-6. Group comparisons made using unpaired T-test. **D.** qRT-PCR analysis of the *MCOLN1* transcripts shows expression of the transgene in whole eye homogenates. Data presented as mean values and SEM; n (HET SALINE)=6; n (KO SALINE)= 5; n (KO scAAV9-JeT-*MCOLN1;* 2e10 vg/mouse)=7; n (KO scAAV9-SYN1-*MCOLN1;* 2e10 vg/mouse)= 4.

Using *MCOLN1* qRT-PCR for biodistribution analysis we found similar expression of the *MCOLN1* transgene in the mouse brain tissue, cortex and cerebellum, as well as in the liver and sciatic nerve **(Figure 5C)**. This is consistent with reports of SYN1 promoter driving expression in hepatocytes and PNS (31). As expected, *MCOLN1* expression was not detected in the muscle tissue of the scAAV9-SYN1-*MCOLN1*-treated *Mcoln1^−/−^* mice **(Figure 5C).** Interestingly, we observed expression of the human *MCOLN1* transgene following scAAV9-Jet-MCOLN1 ICV administration in P1 pups in whole eye homogenates **(Figure 5D)**. This finding warrants further research to establish translational relevance in NHP and human, which would be of high clinical relevance in MLIV.

Additionally, qRT-PCR analysis of the myelination marker *Mbp* showed no change in the *Mbp* transcripts level in the scAAV9-SYN1-*MCOLN1*-treated *Mcoln1^−/−^* mice as compared to the saline-treated *Mcoln1^−/−^* group. Similar to the scAAV9-JeT-*MCOLN1* group, no difference in expression of microglial (*Cd68*) and astrocytic (*Gfap*) markers was observed in scAAV9-SYN1-*MCOLN1*-treated *Mcoln1^−/−^* mice **(Supplementary Figure 3B)**. No significant weight changes or overt health complications were observed in any of the cohorts treated with the scAAV9-MCOLN1 vectors during this study **(Supplementary Figure 3C)**. Overall these data indicate that: 1) CNS, but not muscle, targeting is critical to achieve therapeutic efficacy in MLIV; and 2) within CNS, replacing *MCOLN1* primarily in neurons is sufficient for motor function recovery, at least at the early symptomatic stage of the disease.

### *MCOLN1* gene transfer to peripheral organs fails to rescue disease in *Mcoln1^−/−^* mice

While MLIV primarily affects the CNS, endogenous *MCOLN1* is expressed ubiquitously throughout the body, and the loss off function also impacts peripheral organs, such as muscles, peripheral nerves and stomach (16, 18, 22, 33, 34). We next tested whether systemic administration of scAAV9-JeT-*MCOLN1* in symptomatic mice would have a therapeutic effect in *Mcoln1^−/−^* mice. In this experiment *Mcoln1^−/−^* and control *Mcoln1^+/−^* mice were intravenously administered either saline or the, scAAV9-JeT-*MCOLN1* vector at two months of age **(Table 1)**. Motor function was assessed in the open field test once at the age of 4 months, followed by rotarod testing monthly until the completion of the trial. The qRT-PCR analysis of *MCOLN1* transgene expression showed very low expression in the brain and strong expression in peripheral organs such as skeletal muscle and sciatic nerve, with the highest expression values in the liver **(Figure 6A)**. Despite successful transgene expression in these tissues, we detected no motor function rescue in the scAAV9-JeT-*MCOLN1*-treated *Mcoln1^−/−^* cohort as compared to saline-treated *Mcoln1^−/−^* littermates in either open-field or rotarod tests from 4 to 7 months of age **(Figure 6 B, C, D, E).** This demonstrates that CNS gene transfer is critical to gain therapeutic efficacy. As in all previous trials with *MCOLN1* transfer reported here, no significant weight changes or overt health impacts were noted in the cohort treated intravenously with scAAV9-JeT-*MCOLN1* **(Figure 6F)**.

**Figure 6.**
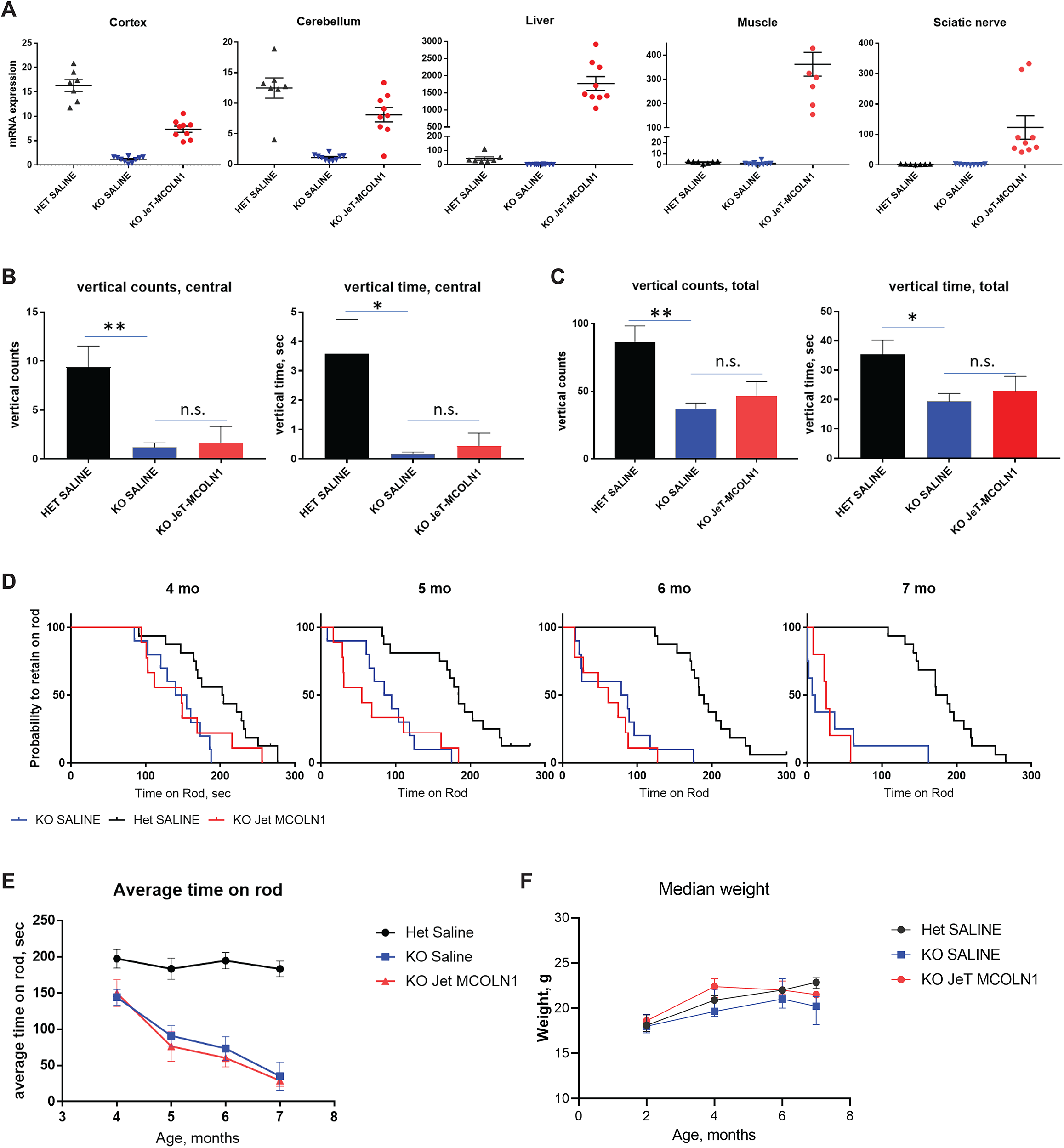
**Brain targeting is required to achieve the therapeutic effect of *MCOLN1* gene transfer. A.** qRT-PCR analysis of the *MCOLN1* transcripts shows that intravenous administration of scAAV9-JeT-*MCOLN1* leads to high expression of *MCOLN1* in peripheral tissues and very low expression in the brain. Data presented as mean values and SEM; n (HET SALINE)=7; n (KO SALINE)= 10; n (KO scAAV9-JeT-*MCOLN1*)=9. **B, C.** Vertical activity deficits are not rescued in *Mcoln1^−/−^* mice after intravenous administration scAAV9-JeT-*MCOLN1*. Vertical activity is presented as vertical counts and vertical time in either central **(B)** or total area **(C)** of the open field arena. N (HET SALINE)=16; n (KO SALINE)=10; n (KO scAAV9-JeT-*MCOLN1*)=9. Data presented as mean values and SEM; group comparisons made using un-paired T-test. **D, E.** Rotarod test data presented either as a probability to retain on the rod **(D)** or average latency to fall **(E)** revealed no improvement in performance of the *Mcoln1^−/−^* mice treated with scAAV9-JeT-*MCOLN1* intravenously. **F.** No significant weight changes have been observed in either experimental group. Data presented as median values and interquartile range, n (HET SALINE) =16; n (KO SALINE) =10; n (KO scAAV9-JeT-*MCOLN1*) =9.

## Discussion

The FDA approval of Zolgensma in 2019 and successful clinical study of an AAV9-based vector for giant axonal neuropathy (GAN) marked a new era for treatment of rare neurological diseases. With more than 200 trials using adeno-associated vectors registered in the US (https://www.clinicaltrials.gov), more than 40 are focused on CNS applications. Several projects, such as in hemophilia A and B, recessive dystrophic epidermolysis bullosa (RDEB), MPSIIIA and MPSIIIB received a “Breakthrough” or “Fast Track” FDA designation with projected approvals by 2022-24. This is changing the landscape of translational research in the rare and ultra-rare disease area and provides a hope for developing transformative therapies.

Based on clinical success, in CNS the vehicle of choice is often a single-stranded or self-complimentary AAV9 vector. AAV9 has high tropism for neurons and other brain cell types, provides long-lasting transgene expression in the brain of rodents, large animals and humans, and shows overall low immunogenicity and toxicity (35, 36). Although AAV9 can cross BBB from systemic flow (37), very high doses are required for systemic delivery to reach therapeutic levels in the CNS in patients older than 2 years of age, prompting the use of direct CNS administration wherever possible. A broad transduction efficiency across the brain and spinal cord can be achieved with intra-CSF administration, either via intracerebroventricular injections in early post-natal period in mice (27), or via intrathecal and intracisternal administration in adult mice (38). Despite having one of the best CNS transduction profiles among naturally occurring AAVs, recombinant AAV9 vectors have limitations in cell type transduction translatability between species (i.e between mice and non-human primates), as well as high pre-existing immunogenicity in the general population that may limit eligibility for treatment (35). Due to technological limitations of the existing vectors, a big effort in the field is now focused on creating novel capsids with improved properties for use in CNS (31, 39–46).

In this study we took advantage of one of such vectors, AAV-PHP.B, which has great properties for CNS transduction from systemic flow 47 and therefor is an excellent tool vector for proof-of-concept experiments targeting the CNS in mice. Using this vector we showed that intravenous delivery in either juvenile (6 weeks) or young adult mice (2 months of age) results in high and long-lasting expression of human transgene in the brain and peripheral organs and, more importantly, it fully restored motor function and delayed time to paralysis in MLIV mice. These data demonstrate that *MCOLN1* gene expression can restore neurologic function in MLIV, even after onset of symptoms.

Since *MCOLN1* product, TRPML1, is ubiquitously expressed and tightly regulated transmembrane channel, our gene transfer strategy for the optimal therapeutic effect was to achieve the broad CNS transduction with moderate expression at the single cell level. Therefore, we used a scAAV9 capsid coupled with the medium strength JeT promotor (26, 29, 48), and ICV route of administration in postnatal day 1 pups. In line with previous reports (31, 42, 48), we detected human *MCOLN1* transgene expression outside of the CNS. Quite surprisingly, we observed low but detectable expression of *MCOLN1* after ICV administration of scAAV9-JeT-MCOLN1 vector in the eye. To our knowledge, this is the first report of the eye transduction following ICV injections. Our data also shows that gene transfer into CNS neurons is critical for functional recovery in MLIV. This is confirmed by the fact that *MCOLN1* expression under the neuron-specific promotor SYN1 resulted in similar recovery of the motor function at the early stage of the disease. Additionally, intravenous administration of scAAV9-JeT-*MCOLN1* in young adult MLIV mice resulting in low *MCOLN1* expression in the brain, failed to rescue neurologic function despite robust expression in peripheral organs, including muscle and peripheral nerves. While the SYN1 and JeT promotor driven constructs resulted in similar functional rescue in *Mcoln1^−/−^* mice, we advocate for use of JeT promoter in therapeutic applications. Given the ubiquitous profile of endogenous *MCOLN1* expression, using the ubiquitous JeT promotor may more accurately recapitulate the reported tissue expression pattern of *MCOLN1*.

In conclusion, we provide experimental evidence that brain-targeted AAV-mediated gene transfer can be a disease altering therapeutic strategy for patients with MLIV. Prompt advancement of gene-therapy trials for rare and devastating disease like MLIV will ultimately help to bring highly efficient gene therapy to the bedside for more common hard-to-treat disorders that seek improved care.

## Materials and Methods

### Animals

*Mcoln1^−/−^* mice were maintained and genotyped as previously described (20). The *Mcoln1^+/−^* breeders for this study were obtained by backcrossing onto a C57Bl6J background for more than 10 generations. Experimental cohorts were obtained from either *Mcoln1^+/−^* × *Mcoln1^+/−^* or *Mcoln1^+/−^* × *Mcoln1^−/−^* mating. *Mcoln1^+/−^* littermates were used as controls. Experiments were performed according to the Institutional and National Institutes of Health guidelines and approved by the Massachusetts General Hospital Institutional Animal Care and Use Committee. Animals were assigned to the experimental groups in a random order and handling and testing was performed by investigator blinded to treatment group info.

### Constructs

Three plasmids were used for *MCOLN1* expression: pAAVss-CMV-*MCOLN1*-FF2A-eGFP, pAAVsc-JeT-*MCOLN1* and pAAVsc-SYN1-*MCOLN1*. The expression vectors were obtained from Dr. Luk Vandenberghe’s lab at Massachusetts Eye and Ear Infirmary (MEEI). pAAVss-CMV-*MCOLN1*-FF2A-eGFP had a bGH polyA, and a human *MCOLN1* cDNA was synthesized and subcloned (https://www.ncbi.nlm.nih.gov/gene/57192). To produce pAAVsc-JeT-*MCOLN1* and pAAVsc-SYN1-*MCOLN1* we used original pAAV SC.CMV.EGFP.BGH vector, where the CMV-eGFP-bGHpolyA expression cassette was replaced by human MCOLN1 cDNA with short synthetic polyA (49) and either minimal synthetic JeT (29) or human SYN1 (50) promoters.

### Virus preparation

AAV-PHP.B-CMV-*MCOLN1*-FF2A-eGFP (referred to as PHP.b-*MCOLN1* in the text) ****and**** scAAV9-JeT *MCOLN1* and scAAV9-SYN1-MCOLN1 viral stocks were prepared at the Gene Transfer Vector Core (http://vector.meei.harvard.edu) at Massachusetts Eye and Ear Infirmary. AAV preparations were produced by triple-plasmid transfection as described previously (45). Near-confluent monolayers of HEK293 cells were used to perform large scale polyethylenimine transfections of AAVcis, AAVtrans, and adeno-virus helper plasmid in a ten-layer hyperflask (Corning, Corning, NY). Downstream purification process, titration and evaluation of purity was performed as described previously (Lock, 2010).

### ICV injections

The viral solution was diluted in sterile saline (0.9% NaCl) with 0.05% trypan blue. Each pup was injected with a maximal injection volume of 5 ul via a 10 ul Hamilton syringe (Hamilton Company, Reno, NV) with a 30G needle. Pups were anesthetized on ice for 5 min until loss of pedal withdrawal reflex. The pup was then placed on a fiber optic light source to illuminate ventricles for injection. The injection site was marked with non-toxic laboratory pen about 0.25 mm lateral to sagittal suture and 0.50-0.75 mm rostral to neonatal coronary suture (0.25 mm lateral to confluence of sinuses). Pups were injected with either saline, scAAV9-JeT-*MCOLN1* at a dose of 2 × 10^10^ vg/mouse, 1 × 10^10^ vg/mouse or 0.4 × 10^10^ vg/mouse or scAAV9-SYN1-*MCOLN1* at a dose of 2 × 10^10^ vg/mouse. After, injection the pup was placed under a warming lamp and monitored for 5-10 min for recovery until movement and responsiveness was fully restored.

### IV injections

Mice were restrained in a plexiglass restrainer. The tail veins were dilated in warm water for 1 minute. 30G-needle insulin syringes (Cat. No 328466,BD, San Diego, CA) were used with total injection volume of 70-100 ul per mouse. Mice were injected with either saline, AAV-PHP.b-CMV-*MCOLN1* at a dose of 1 ×10^12^ vg/mouse or scAAV9-JeT-*MCOLN1* at a dose of 5 × 10^11^ vg/mouse.

### Behavioral testing

Open field testing was performed on naive male and female mice at either two or four months of age under regular light conditions. Each mouse was placed in the center of a 27 × 27 cm^2^ Plexiglas arena, and the horizontal and vertical activity were recorded by the Activity Monitor program (Med Associates Fairfax, VT). Data were analyzed during the first 15 minutes in the arena. Zone analysis was performed to, measure movements/time spent in the central (8 × 8 cm^2^) versus peripheral (residual) zone of the arena. Statistical significance was determined using an unpaired T-test.

Motor coordination and balance was tested in on an accelerating rotarod (Med Associates, VT). Latency to fall from the rotating rod was recorded in 3 trials per day (accelerating speed from 4 to 40 rpm over 5 min) during 2 days of testing. To follow the progression of motor decline, performance of mice was tested monthly starting at 4 months until the motor dysfunction in *Mcoln1^−/−^* mice made it impossible to perform the task. Cox-regression and log-rank analysis was used to determine probability of falling from the rod.

### RNA Extraction and qPCR Analysis

Mouse tissues were disrupted and homogenized using QIAzol lysis reagent (Qiagen, Hilden, Germany) and the Tissue Lyser instrument (Qiagen, Hilden, Germany). 30 mg of snap frozen tissues were processed adding 1 ml of QIAzol reagent in presence of one 5 mm stainless steel bead (Qiagen, Hilden, Germany). Total RNA isolation from homogenized tissues was performed using Qiagen RNeasy kit (Qiagen, Hilden, Germany) and genomic DNA was eliminated performing DNase (Qiagen, Hilden, Germany) digestion on columns following procedures indicated by the provider. cDNA was generated using High-Capacity cDNA Reverse Transcription kit (Applied Biosystems, Foster City, CA) according to manual instruction. cDNA produced from 500 ng of starting RNA was diluted and 40ng were used to perform qPCR using LightCycler 480 Probes Master mix (Roche Diagnostics, Mannheim, Germany). The real-time PCR reaction was run on LightCycler 480 (Roche Diagnostics, Mannheim, Germany) using TaqMan premade gene expression assays (Applied Biosystems, Foster City,CA). Mus musculus: GAPDH (FAM)-Mm99999915_g1, MBP, (FAM)- Mm01266402_m1, GFAP (FAM)-Mm01253033_m1, CD68 (FAM)-Mm03047343_m1; Homo Sapiens

Mucolipin-1 (FAM)-Hs01100653_m1. The ΔΔCt method was used to calculate relative gene expression, where Ct corresponds to the cycle threshold. ΔCt values were calculated as the difference between Ct values from the target gene and the housekeeping gene GAPDH.

### Tissue collection and processing

Mice were sacrificed using a carbon dioxide chamber. Immediately after euthanasia, mice were transcardially perfused with ice-cold phosphate buffered saline (PBS). After bisecting the brain across the midline, half was used to isolate cerebellum and cortex. The other half was post-fixed in 4% paraformaldehyde in PBS for 48 h, washed with PBS, cryoprotected in 30% sucrose in PBS for 24 h, frozen in isopentane and stored at −80°C. In addition to brain tissue, liver, muscle and sciatic nerve tissue were collected and snap-frozen over dry ice before storage at −80°C.

### Immunohistochemistry, imaging, and analysis

For histological analysis, 40 μm coronal sections were cut using a cryostat (Leica Microsystems, Wetzlar, Germany) and collected into 96 well plates containing cryoprotectant (TBS, 30% ethylene glycol, 15% sucrose). These sections were stored at 4°C prior or frozen at -80C. Prior to staining for LAMP1 and Myelin Basic Protein (MBP), samples were randomized and coded to create blinded conditions for analysis. Staining of free-floating sections was done in a 96 well plate. Sections were blocked in 0.1% Triton x-100,10% normal horse serum (NHS), 2% bovine serum albumin (BSA), and 1% glycine in PBS and incubated with primary antibodies diluted in antibody buffer: 10% NHS and 2% BSA overnight. The following primary antibodies were used: a-LAMP1 antibody (Rat 1:1000, BD San Diego, CA, Cat ID: 553792); MBP (Mouse,1:1000, Millipore Sigma, Jaffrey, NH, NE1019). Sections were washed in PBS-Tween (0.05%) at room temperature then incubated with secondary antibodies in antibody buffers for 1 hour at room temperature. The following secondary antibodies were used: goat-anti-rat AlexaFluor 546 (1:500; Invitrogen, Eugene, OR), goat-anti-mouse AlexaFluor 555 (1:500; Invitrogen, Eugene, OR). Sections were counterstained with NucBlue nuclear stain (Life Technologies, Eugene, OR). After staining, sections were mounted onto glass SuperFrost plus slides (Fisher Scientific, Pittsburgh, PA) with Immu-Mount (Fisher Scientific, Pittsburgh, PA).

Images were acquired on DM8i Leica Inverted Epifluorescence Microscope with Adaptive Focus (Leica Microsystems, Buffalo Grove, IL) with Hamamatsu Flash 4.0 camera and advanced acquisition software, package MetaMorph 4.2 (Molecular Devices, LLC, San Jose, CA) using an automated stitching function. The exposure time was kept constant for all sections within the same IHC experiment. Image analysis was performed using Fiji software (NIH, Bethesda, Maryland). For corpus callosum area measurements, areas of interest were selected on MBP images in each section and normalized to the whole section area. For LAMP1 images, particle analysis and percent of area measurements were made after same thresholding settings were allied to all images. All images were decoded after the measurements were taken. Area and mean pixel intensity values were averaged per genotype/treatment group and compared between genotypes using unpaired T-test.

### Statistical analysis

Data presented as mean values and SEM or median values and interquartile range. Statistical analysis was performed in GraphPad Prism 7.04 software (GraphPad, La Jolla, CA) using either unpaired T-test, two-way ANOVA or log-rank as tests detailed in specific subsections of Methods. P values were indicated as follows throughout the manuscript: n.s. = p < 0.05; * p =< 0.05; ** p =< 0.01, *** p = < 0.001,**** p =< 0.0001.

## Acknowledgments

This work has been funded by the grant to YG from the ML4 and Zalik Foundations. Authors are especially grateful to Dr. Rebecca Oberman and Randy and Caroline Gold for their long-term support of the MLIV research program at Mass General Research Institute and dedication and help to the MLIV community. YG and SAS are inventors on the IP filed through Mass General Brigham Corporation. Authors are grateful to Drs. Florian Eichler and Tim Riley for fruitful discussions.

## Author Contributions

S.D.- Investigation, Writing – original draft; M.S.- Investigation, Methodology, Supervision, Writing – review & editing; S.S. – Investigation, Methodology, M.S. – Investigation; V.M.-B. - Writing – review & editing, Investigation; S. W.- Supervision, Writing – review & editing; R. X.- Methodology, L. V. – Conceptualization; S. S.- Writing – review & editing; A. M. - Writing – review & editing; Y. G.- Conceptualization, Formal Analysis, Funding acquisition, Investigation, Visualization, Writing – original draft.

---

**Supplementary Figure 1. Open field test reveals robust motor deficits in *Mcoln1^−/−^* mice.** Data of the first 15 min of exploration in the arena are shown. Male and female *Mcoln1^−/−^* mice demonstrate significantly reduced vertical activity presented in the form of vertical counts and vertical time in the center of the arena and in the total area of the arena. N, males: (WT, 1 mo)=13, (KO, 1mo)=9, (WT, 2mo) = 17, (KO, 2 mo) = 17, (WT, 4mo) = 14, (KO, 4mo) = 9, (WT, 5mo) =7, (KO, 5mo)=8; females: (WT, 1mo)=8, (KO, 1mo) = 8, (WT, 2mo) = 22, (KO, 2mo) = 36, (WT, 4mo)=28, (KO, 4mo) =28, (WT, 5mo) = 10, (KO, 5mo) = 8.

**Supplementary Figure 2. Additional timepoints of the rotarod testing at 8 and 9 months of age in *Mcoln1^−/−^* (KO PHP.b-*MCOLN1*) and control mice (HET SALINE) treated at symptomatic stage of the disease at 2 months, related to Figure 3.**

**Supplementary Figure 3. Intracerebroventricular delivery of scAAV9-JeT-*MCOLN1* in P1 *Mcoln1^−/−^* doesn’t affect glial activation or myelination marker *Mbp* expression in the brain. A.** qRT-PCR analysis of the microgliosis marker *Cd68*, astrocytosis marker *Gfap* and myelination marker *Mbp* in the cerebral cortex shows no correction in *Mcoln1^−/−^* mice regardless scAAV9-JeT-*MCOLN1* dose. Data presented as mean values and SEM; n (HET SALINE)=20; n (KO SALINE)= 10; n (KO scAAV9-JeT-*MCOLN1;* 2e10 vg/mouse)=7; n (KO scAAV9-JeT-*MCOLN1;* 1e10 vg/mouse)= 5; n (KO scAAV9-JeT-*MCOLN1;* 4e9 vg/mouse)= 10. **B.** qRT-PCR analysis of the microgliosis marker *Cd68*, astrocytosis marker *Gfap* and myelination marker *Mbp* in the cerebral cortex following ICV administration at P1 of either scAAV9-JeT-*MCOLN1* or scAAV9-SYN1-*MCOLN1* vectors. Data presented as mean values and SEM; n (HET SALINE)=4-5; n (KO SALINE)= 4-5; n (KO scAAV9-JeT-*MCOLN1;* 2e10 vg/mouse)=6; n (KO scAAV9-SYN1-*MCOLN1;* 2e10 vg/mouse)= 4-6. Group comparisons made using unpaired T-test. **C.** No significant weight changes have been observed in either experimental group following ICV injections at P1 with scAAV9 *MCOLN1* expressing vectors. Data shows only male mice in the experimental cohort and presented as median values and interquartile range, n (HET SALINE) =17; n (KO SALINE) = 10; n (KO scAAV9-JeT-*MCOLN1;* 2e10 vg/mouse)=7; n (KO scAAV9-JeT-*MCOLN1;* 1e10 vg/mouse)= 4; n (KO scAAV9-JeT-*MCOLN1;* 4e9 vg/mouse)= 10; n (KO scAAV9-SYN1-*MCOLN1;* 2e10 vg/mouse)= 6.

