## Supplementary figures and images for "*MCOLN1* gene-replacement therapy corrects neurologic dysfunction in the mouse model of mucolipidosis IV"

### Supplementary Figure 1

Supplementary figure 1

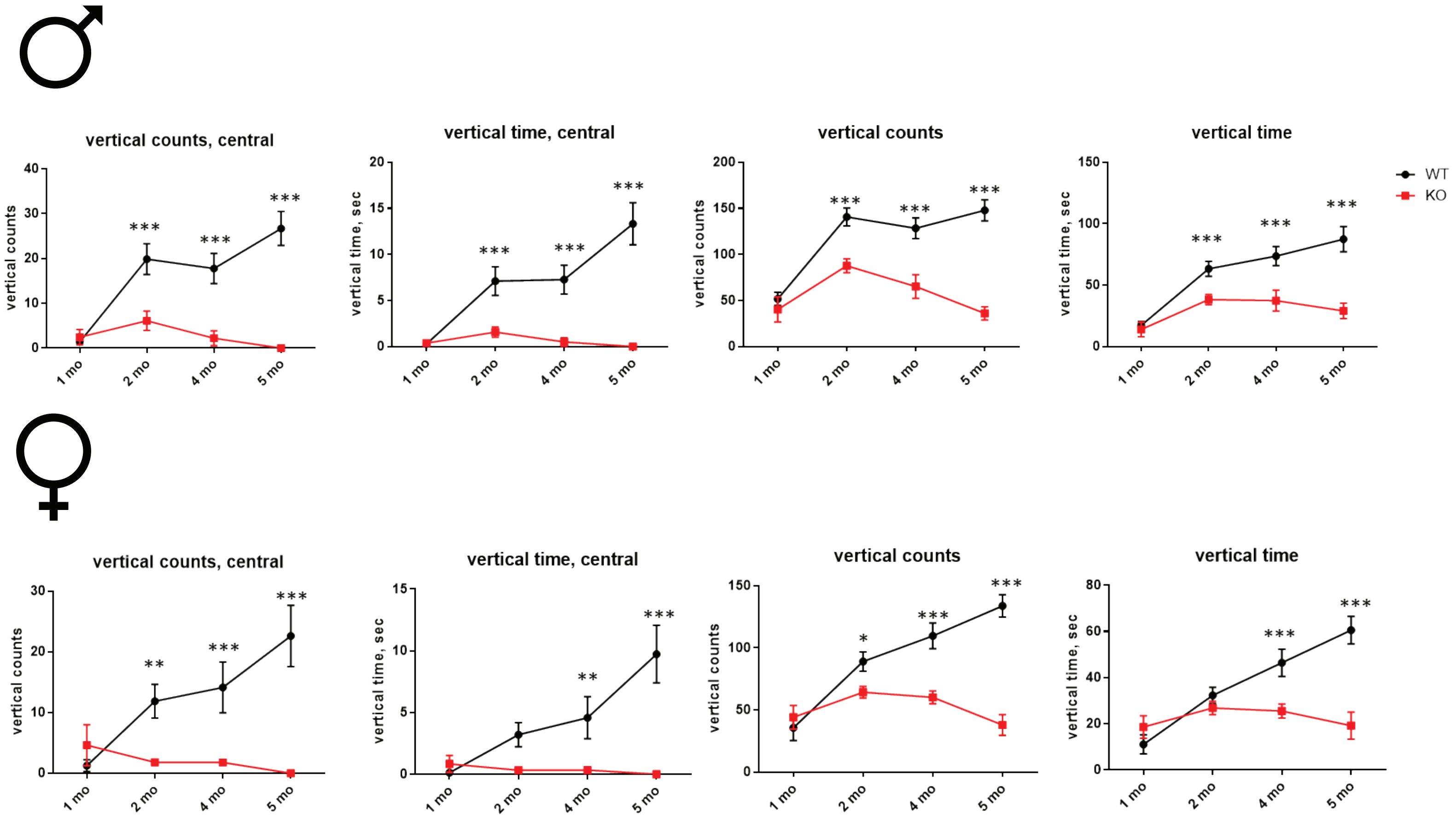

### Supplementary Figure 2

Supplementary figure 2

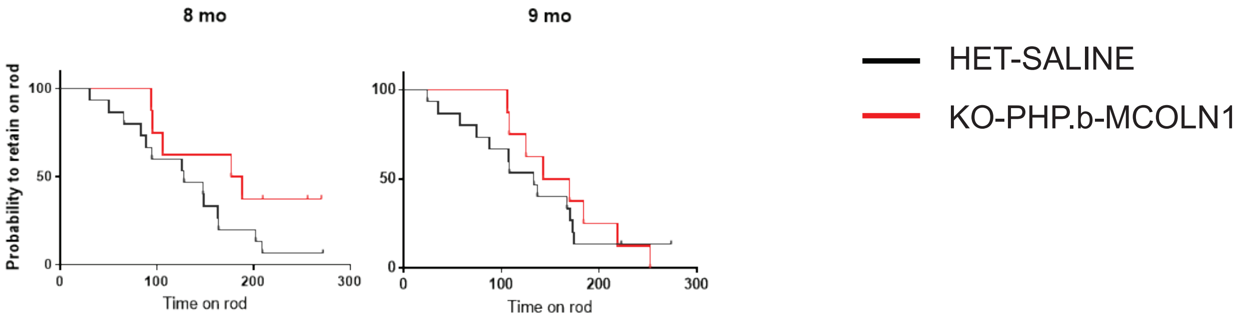

### Supplementary Figure 3

Supplementary figure 3

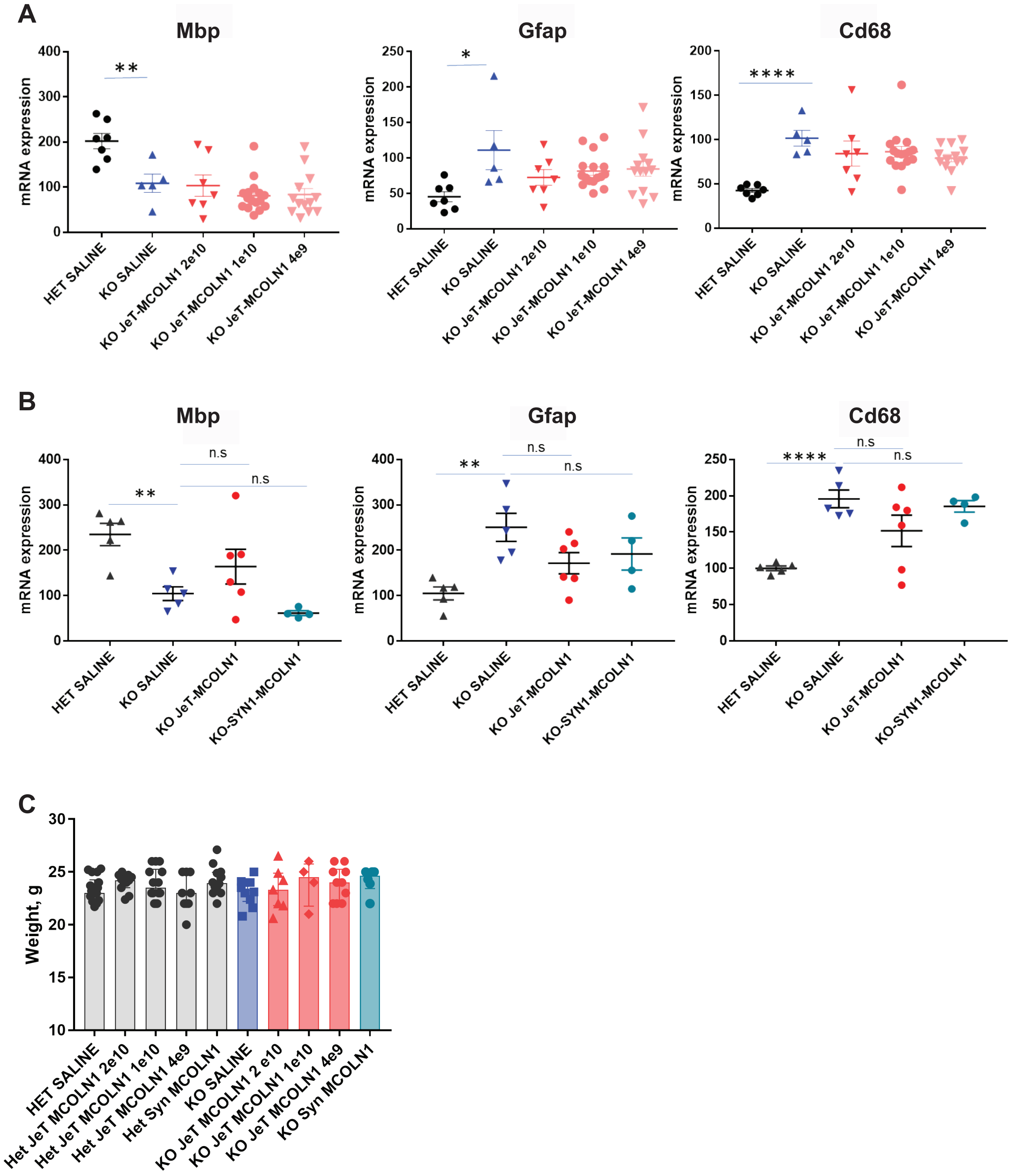
